# A Genetic Screen Identifies Two Novel Rice Cysteine-rich Receptor-like Kinases That Are Required for the Rice NH1-mediated Immune Response

**DOI:** 10.1101/003129

**Authors:** Mawsheng Chern, Rebecca S. Bart, Wei Bai, Deling Ruan, Wing Hoi Sze-To, Patrick E. Canlas, Rashmi Jain, Xuewei Chen, Pamela Ronald

**Affiliations:** Department of Plant Pathology and the Genome Center, University of California, Davis. Davis, CA 95616. USA; Joint Bioenergy Institute, Emeryville, California, USA; College of Life Sciences, Inner Mongolia Agricultural University, Huhhot 010018, China; Rice Research Institute, Sichuan Agricultural University at Chengdu, 211 Huimin Road, Wenjiang, Chengdu, Sichuan, 611130, China; Current Address: Donald Danforth Center, 975 North Warson Rd. St. Louis, MO 63132. USA

## Abstract

**Abbreviations**
BTH: benzothiadiazole
CRK: cysteine-rich receptor-like kinases
NH1: NPR1 homolog 1
NPR1: non-expressor of pathogenesis-related genes 1
SA: salicylic acid
SAR: systemic acquired resistance; *sn11*, suppressor of NH1-mediated lesion mimics 1.

Over-expression of rice *NH1* (NH1ox), the ortholog of Arabidopsis *NPR1*, confers immunity to bacterial and fungal pathogens and induces the appearance of necrotic lesions due to activation of defense genes at the pre-flowering stage. This lesion-mimic phenotype can be enhanced by the application of benzothiadiazole (BTH). To identify genes regulating these responses, we screened a fast neutron-irradiated NH1ox rice population. We identified one mutant, called *sn11* (<u>s</u>uppressor of <u>N</u>H1-mediated lesion-mimic 1), which is impaired both in BTH-induced necrotic lesion formation and in the immune response. Using a comparative genome hybridization approach employing rice whole genome tiling array, we identified 11 genes associated with the *sn11* phenotype. Transgenic analysis revealed that RNA interference of two of the genes, encoding previously uncharacterized cysteine-rich receptor-like kinases (CRK6 and CRK10), re-created the *sn11* phenotype. Elevated expression of *CRK10* using an inducible expression system resulted in enhanced immunity. Quantitative PCR revealed that BTH treatment and elevated levels of rice NH1 and its paralog NH3 induced expression of *CRK10* and *CRK6* RNA. These results indicate that CRK6 and CRK10 are required for the BTH-activated immune response mediated by NH1.

## Introduction

Plants survive pathogen attack by employing various defense strategies, including strengthening their cell walls, generation of reactive oxygen species, accumulating phytoalexins, and synthesizing salicylic acid (SA) [1]. After initial local infection, most plants are able to initiate a defense response termed systemic acquired resistance (SAR), which includes induction of expression of a set of pathogenesis-related (*PR*) genes, leading to a long-lasting enhanced resistance against a broad spectrum of pathogens [2]. In dicots, SA and its synthetic analogs, 2,6-dichloroisonicotinic acid (INA), benzothiadiazole (BTH), and probenazole, are potent inducers of SAR [3–5]. Methyl salicylate, rather than SA, is the critical mobile signal for SAR [6,7]. In wheat, SAR is induced by BTH treatment [8] and in rice, by *Pseudomonas syringae* [9]. BTH also induces disease resistance in rice [10–12] and maize [13], although it is unclear if these defense responses are equivalent to SAR.

The *NPR1* (nonexpressor of pathogenesis-related genes 1; also known as *NIM1* and *SAI1*) gene is a key regulator of SA-mediated SAR in Arabidopsis [14–18]. Upon induction by SA, INA, or BTH, *NPR1* expression levels increase, influencing the SAR response [19]. Arabidopsis *npr1* mutants are impaired in their ability to induce *PR* gene expression and cannot mount a SAR response even after treatment with SA or INA. In Arabidopsis, over-expression of *NPR1* leads to enhanced resistance to both bacterial and oomycete pathogens [20,21]. *NPR1* encodes a protein with a bipartite nuclear localization sequence and two protein-protein interaction domains: an ankyrin repeat domain and a BTB/POZ domain [19].

Research over the last decade has partially revealed the mechanism of action of this important regulator. NPR1 forms an oligomer before activation that is mostly excluded from the nucleus. Upon SAR induction and subsequent change to the cellular redox state, monomeric NPR1 is released and accumulates in the nucleus, activating *PR* gene expression [22]. NPR1 interacts with TGA transcription factors [23–25], which mediate its function [26,27]. NPR1 functions as a transcriptional co-activator in a TGA2-NPR1 complex after SA treatment in a transient cell assay; this function requires the BTB/POZ domain and the oxidation of NPR1 Cys-521 and Cys-529 [28]. The NPR1 BTB/POZ domain interacts with the repression domain of TGA2 to neutralize its repression function [29]. The BTB/POZ domain also serves to sequester and repress the C-terminal transactivation domain of NPR1 and SA induction may release this inhibition [30].

It has been hypothesized that Arabidopsis NPR1 is an SA receptor that directly binds to SA, resulting in a conformational change that releases its C-terminal transcriptional activation domain and transforms the NPR1 protein into a functional transcriptional co-activator [30]. Another report demonstrates that Arabidopsis NPR3 and NPR4 have a higher binding affinity for SA than NPR1. In this model, NPR3 and NPR4 are the SA receptors. Binding of SA to NPR3 or NPR4 triggers NPR1 degradation mediated by the Cullin 3 ubiquitin E3 ligase [31]. Both these models indicate that SA modulates NPR1 function.

In rice, over-expression of Arabidopsis *NPR1* [25] or the rice ortholog *NH1* [32,33] results in enhanced resistance to the pathogens *Xanthomonas oryzae* pv. *oryzae* (*Xoo*) and *Magnaporthe grisea*, the causal agents of rice bacterial leaf blight and rice blast, two of the most destructive rice diseases worldwide. Of the five rice NPR1-like genes in rice, only *NH1* and *NH3* enhance resistance to *Xoo* when expressed at elevated levels [33,34]. Rice NH1 also interacts with TGA transcription factors [32]. The enhanced disease resistance of NH1ox rice plants is accompanied by cell death, commonly referred to as a lesion mimic phenotype [32,35,36]. The development of lesion mimic necrotic spots at the pre-flowering stage correlates with enhanced resistance to *Xoo* and induction of *PR* gene expression [32]. Application of BTH to the NH1ox plants greatly enhances the formation of necrotic spots, suggesting that the NH1-mediated lesion mimic phenotype is tightly associated with BTH-induced resistance in rice.

Although Arabidopsis NPR1 and rice NH1 have been shown to act as transcriptional co-activators [28,37], activating target genes by binding to transcription factors, such as TGA proteins, the downstream components that mediate the signaling cascade leading to the immune response remain uncharacterized. To identify such proteins, we initiated a suppressor screen to identify genes involved in the BTH-induced, NH1-mediated immune response. We generated a rice mutant population by treating NH1ox rice seeds with fast-neutron radiation, which generally leads to small deletions [38] and substitutions [39]. In a screen of approximately 60,000 M2 treated with BTH, we identified <u>s</u>uppressors of <u>N</u>H1-mediated <u>l</u>esion-mimic (*snl*) mutants that no longer respond to BTH. Here, we report the characterization of a mutant, *snl1,* and the isolation of two previously uncharacterized cysteine-rich receptor-like kinases (*CRK6* and *CRK10*) that are required for the BTH-induced immune response.

## Results

### Identification of mutant *snl1* and the deletion associated with the suppressor phenotype by comparative genomic hybridization

In our mutant screen for suppressors of BTH-induced, NH1-mediated immunity, we identified multiple *snl* mutants [40]. Here, we report the characterization of mutant *snl1*. As shown in Figure 1A, the *snl1* mutant displayed no necrotic spots after application of the chemical inducer BTH, while the NH1ox parent displayed typical lesion mimic necrotic spots. After inoculation with *Xoo* strain PXO99, the *snl1* mutant developed long water-soaked lesions typical of the disease whereas the NH1ox parental line developed much shorter disease lesions, as shown in Figure 1B. The difference in lesion lengths was confirmed by measuring bacterial populations in leaves of *snl1* and NH1ox plants. Figure 1C shows that *snl1* harbored an 18-fold larger *Xoo* population compared to the NH1ox control with a P value of 0.0052 on T-test. These results confirm that the *snl1* mutant is compromised in resistance to *Xoo* and NH1ox-mediated lesion mimic development.

**Figure 1.**
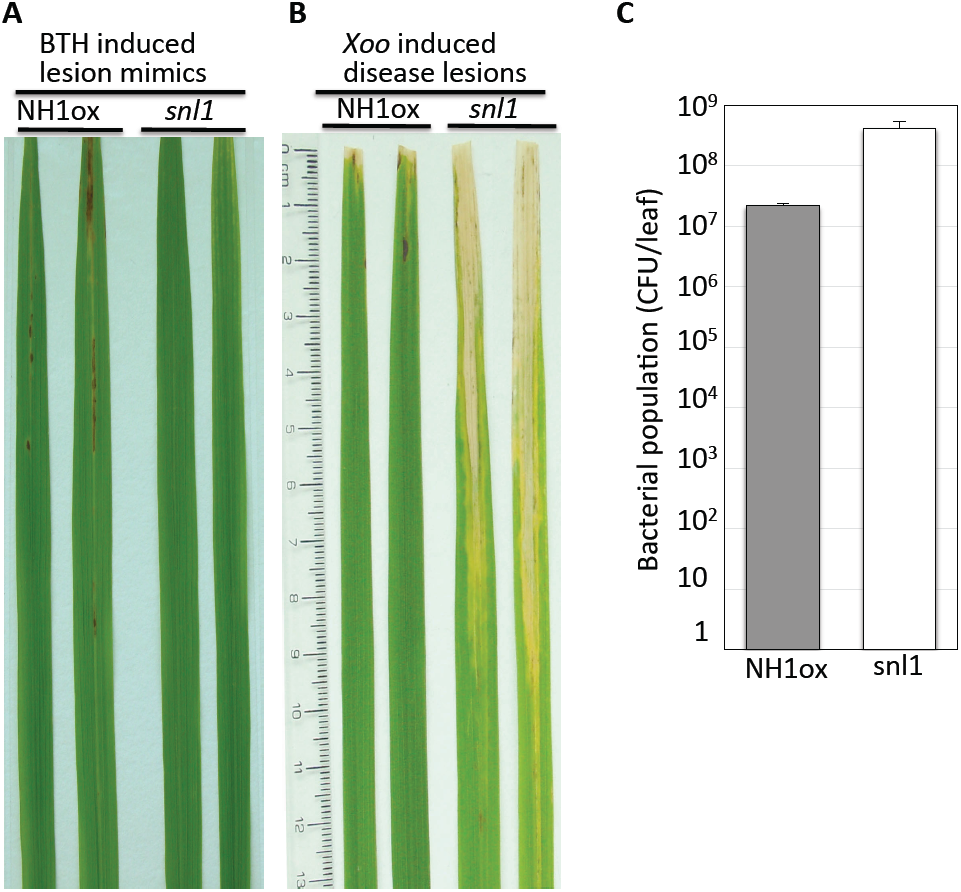
The *snl1* mutant is compromised in BTH-induced necrotic lesion formation and resistance to *Xoo*. Two representative leaves are displayed for each of the NH1ox parent and the *snl1* mutant in (A) and (B). Inoculation with PXO99 was carried out with the scissor-dip method described previously (see Methods). (A) Lesion mimic necrotic spots. (B) *Xoo*-induced, water-soaked disease lesions and (C) Bacterial populations 14 days after inoculation. T-test yielded P=0.0052. Each bar represents the mean and standard deviation of three leaves.

To expedite the isolation of the gene(s) responsible for the *snl1* mutation, we carried out CGH analysis on *snl1* genomic DNA and that of its parent (NH1ox) using a NimbleGen 2.1-million probe rice whole genome tiling array, which carries on average one probe per 150 bp and has higher probe density in genic regions [40]. A summary of the CGH results for mutant *snl1* is shown in Figure 2A, where a downward peak represents a deletion on the 12 chromosomes, depicted by the 12 different colors. A single large (∼88 kb) deletion is present on chromosome 7 in mutant *snl1* as shown in Figure 2B, where each dot represents a probe on the microarray. This single 88-kb deletion in *snl1* was confirmed by PCRs comparing *snl1* genomic DNA with that of NH1ox, targeting 9 genes in this region (data not shown).

**Figure 2.**
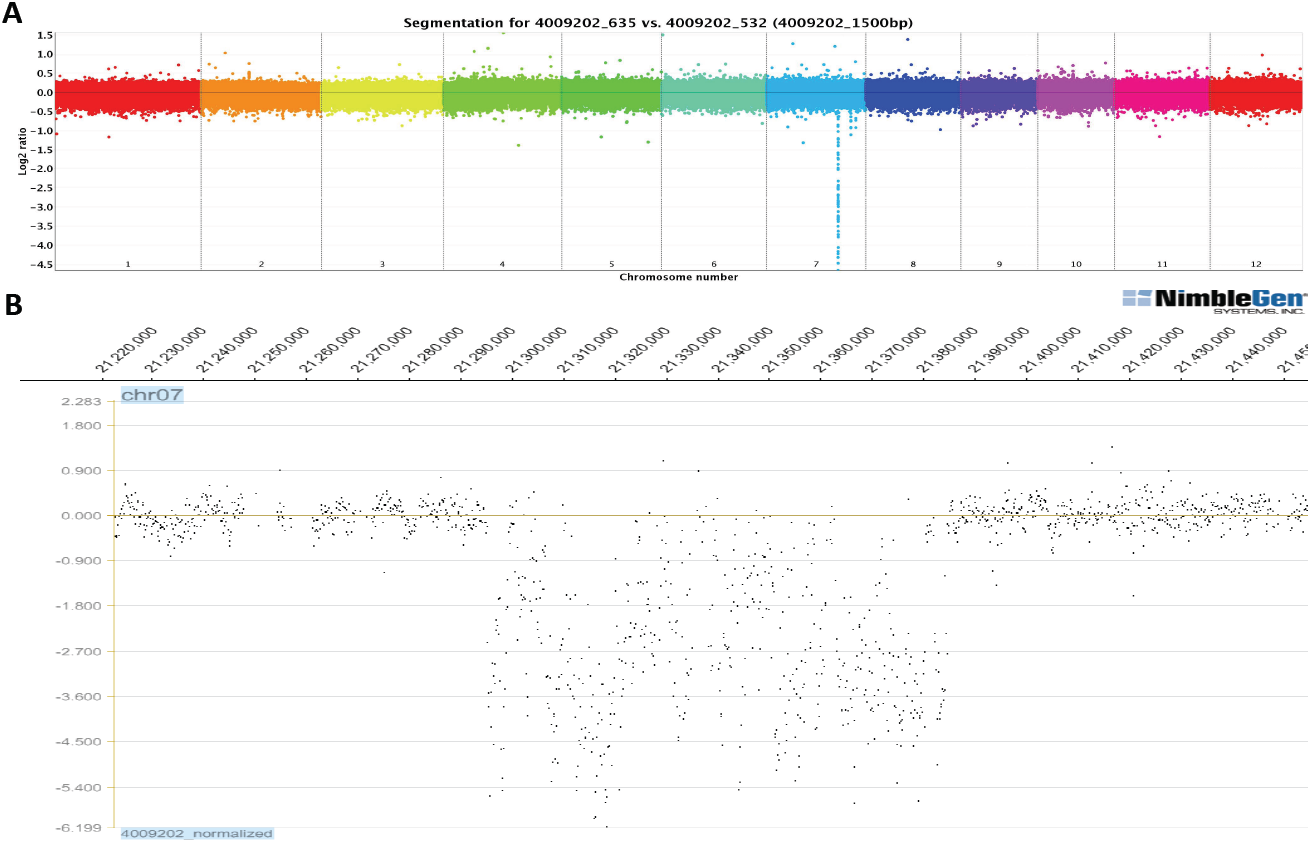
Comparative genome hybridization identifies an 88-kb deletion on chromosome 7 of *snl1*. Purified genomic DNA samples of the *snl1* mutant and its NH1ox parent were labeled with cy3 and cy5 separately and hybridized to NimbleGen 2.1-M element rice tiling array. (A) A composite graph of the 12 rice chromosomes. Each color represents a chromosome; each dot represents a 1-kb region on chromosome. The downward peak on chromosome 7 represents a deletion. (B) Actual hybridization data of *snl1* on a segment of chromosome 7. Each dot represents an actual probe on the array. The signals derived from probes are represented in numbers of log2. The downward shifts of signals suggest a large deletion in mutant *snl1* that was confirmed by PCR genotyping.

The Nipponbare reference genome shows 11 annotated genes in the *snl1*deleted region: 6 cysteine-rich receptor-like kinases (CRKs, including MSU locus ID Os07g35580, Os07g35650, Os07g35660, Os07g35680, Os07g35690, and Os07g35700), one DCD (development and cell death)-kelch motif protein (Os07g35610), two expressed proteins (Os07g35600 and Os07g35630), and 2 hypothetical proteins (Os07g35620 and Os07g35640).

### The 88-kb deletion cosegregates with the *snl1* phenotype

The *snl1* mutant was crossed with the Liaogeng (LG) parent, from which the NH1ox line was generated via introducing the *Ubi-NH1* gene. An F2 segregating population derived from the F1 was analyzed for susceptibility to *Xoo* and scored for the presence of the *Ubi-NH1* gene and the 88-kb deletion, represented by genes Os07g35610, Os07g35690 (encoding CRK6), and Os07g35700 (encoding CRK10). Our PCR analysis results revealed that Os07g35610, *CRK6*, and *CRK10* were deleted in the mutant segregants. Among the progeny containing the *Ubi-NH1* gene, those that still contained the *CRK10* gene (labeled as *CRK10* positive, or group A) all exhibited resistance to *Xoo* at levels similar to the NH1ox parent, albeit with some variations (Figure 3). Progeny lacking the *CRK10* gene (*CRK10* negative, or group B) all displayed susceptibility to *Xoo*, similar to progeny lacking the *Ubi-NH1* gene. Thus, the deletion of *CRK10* is completely associated with the *snl1* phenotype. Plants that lack the *Ubi-NH1* gene display susceptibility to *Xoo* [25].

**Figure 3.**
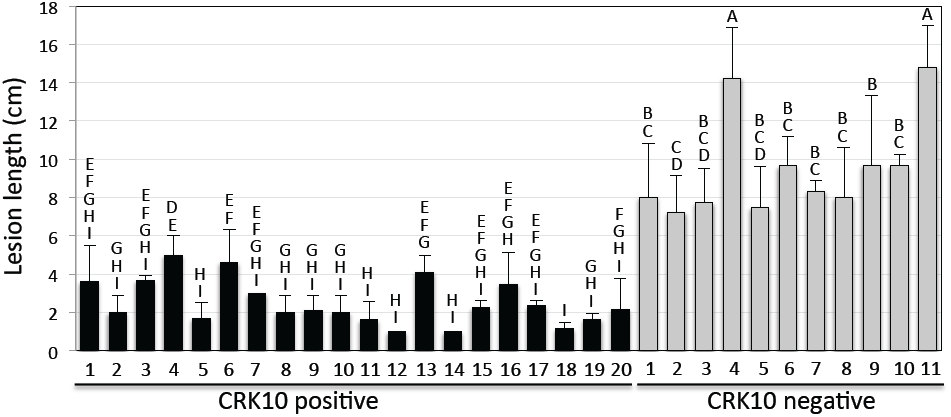
The 88-kb deletion cosegregates with the *snl1* phenotype. A segregating progeny population derived from a cross between the *snl1* mutant and the LG parent was genotyped for the presence of the *Ubi-NH1* gene (NH1ox) and the *CRK10* gene (representing the 88-kb deletion) and scored for resistance to *Xoo*. Only progeny containing the *Ubi-NH1* gene were presented for the phenotype. *Xoo* caused lesion lengths were measured and plotted for the progeny. Those progeny containing the *CRK10* gene are presented as filled bars and those missing the *CRK10* gene presented as grey bars. The letters above each bar show the statistical groupings using the student T-test on each pair based on the 5% significance level.

T-test revealed that the difference between group A (*CRK10* positive) and group B (*CRK10* negative) is highly significant with a p value lower than 0.0001. This experiment was performed twice and similar results were obtained each time. The variations observed among Group A might represent the difference in the copy number of *Ubi-NH1* and *CRK10*. These results clearly demonstrate that the susceptible phenotype of *snl1* is tightly associated with the 88-kb deletion.

### The *snl1* mutation compromises BTH-induced resistance to *Xoo* independent of ectopic NH1 over-expression

In order to assess the role of the *snl1* mutant under physiological conditions, we tested if this mutation affects resistance to *Xoo* in a genetic background where the *NH1* gene is not ectopically over-expressed. For this purpose, we identified a progeny line (line # 43, abbreviated as *snl1*/LG) in which the *Ubi-NH1* gene is absent and the *snl1* 88-kb deletion is homozygous from the cross of *snl1*xLG described above. We tested 27 progeny (110 leaves) of this *snl1*/LG line, together with 27 plants (93 leaves) of the LG parental control. These plants were treated with 1 mM BTH and inoculated with PXO99 two days later (Figure 4). The *snl1*/LG line displayed an average lesion length of 9.0±1.9 cm whereas the wild type LG had an average of 6.0±1.5 cm. T-test on the two sets of lesion length data yielded a P value lower than 0.0001, indicating a highly significant difference between the lesion lengths of the two populations. These results confirm that the *snl1* mutation affects BTH-induced resistance to *Xoo* in a wild type rice genetic background.

**Figure 4.**
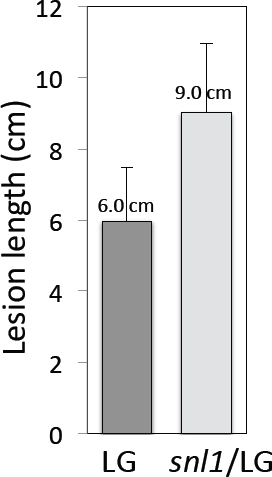
The *snl1* mutation affects resistance to *Xoo* in the wild type genetic background. A population of 27 plants of snl1/LG was inoculated with PXO99 together with 27 plants of the LG control. Lesion lengths were measured 14 days after inoculation. The average lesion lengths and standard deviations were presented in the graph. T-test between LG and snl1/LG gave a P value lower than 0.0001.

### *CRK6* and *CRK10* partially complement the suppressor phenotype of *snl1*

In order to assess which gene(s) in the 88-kb deletion is responsible for the *snl1* phenotype, we carried out complementation experiments for each of the 11 annotated genes in the region. For complementation, we used long-range PCR to amplify each gene. Each gene, including the coding region, the putative promoter (approximately 1.5 kb upstream of the start codon), and the 3’ untranslated and untranscribed regions (approximately 500 bp after the stop codon), was cloned and confirmed by sequencing. Each gene was then cloned into the C4300 binary vector (see Methods).

Transformation of mutant *snl1* (hygromycin resistant) with each of these constructs was carried out using mannose selection. We generated more than 20 independent transgenic lines each for Os07g35630, Os07g35660, Os07g35680, *CRK6* and *CRK10*, more than 10 lines for Os07g35610, Os07g35620, and Os07g35650, 5 lines for Os07g35640 and Os07g35580, and 4 lines for Os07g35600.

We challenged the T0 transgenic plants with *Xoo* strain PXO99 and measured water-soaked lesions 14 days after infection. We found that none of the individual genes were able to fully restore resistance to *Xoo* and lesion mimic development to the levels of the NH1ox parent. However, most of the CRK6 and CRK10 transgenic lines were more resistant to *Xoo* than the *snl1* parent, displaying shorter lesion lengths, as shown in Figure 5. The fact that both *CRK6* and *CRK10* are capable of partially complementing the *snl1* mutant suggests that CRK6 and CRK10 might contribute to the *snl1* phenotype.

**Figure 5.**
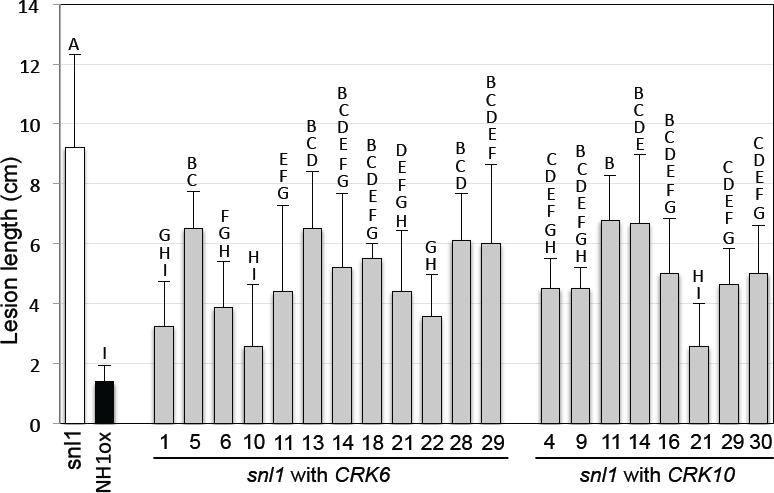
*CRK6* and *CRK10* each partially complements the *snl1* phenotype. The *snl1* mutant was complemented with *CRK6* or *CRK10*. The complementation lines and the *snl1* and NH1ox controls were inoculated with PXO99 and lesions measured 14 days after infection. Twelve independent *CRK6* and eight *CRK10* complementing lines are shown. Each bar represents the average and standard deviation of at least three leaves. The letters above each bar show the statistical groupings using the student T-test on each pair based on the 5% significance level.

### Expression levels of *CRK6* and *CRK10* are induced by BTH and elevated in *NH1* or *NH3* over-expression plants

Using real time quantitative reverse transcription (RT)-PCR, we tested *CRK6* and *CRK10* expression levels after treatment with 1 mM BTH in wild type Kitaake (Kit) and nNH1 and nNH3 transgenic plants, which express higher levels of *NH1* and *NH3* (driven by their native promoters in Kit background) [34], respectively. As shown in Figure 6A, *CRK6* expression was induced 6 fold (peaked at 4 hours after induction) in wild type Kit. *CRK6* induction by BTH was slightly delayed in nNH1 plants (peaked at 8 hours instead of 4 hours). The *CRK6* expression level was elevated in nNH3 plants before BTH induction and showed similar induction pattern as in Kit after BTH treatment. The results in Figure 6B show that the *CRK10* level was induced nearly 3-fold (peaked at 8 hours after induction) in Kit. *CRK10* expression was elevated 2.5-fold before BTH induction in nNH1 plants, and was further induced by BTH treatment to 4.5-fold of the uninduced level in Kit. *CRK10* expression showed little difference in nNH3 and Kit with or without BTH induction. These results indicate that *CRK6* and *CRK10* expression are both induced by BTH treatment, that NH3 induces *CRK6* expression, and that *NH1* induces *CRK10* expression.

**Figure 6.**
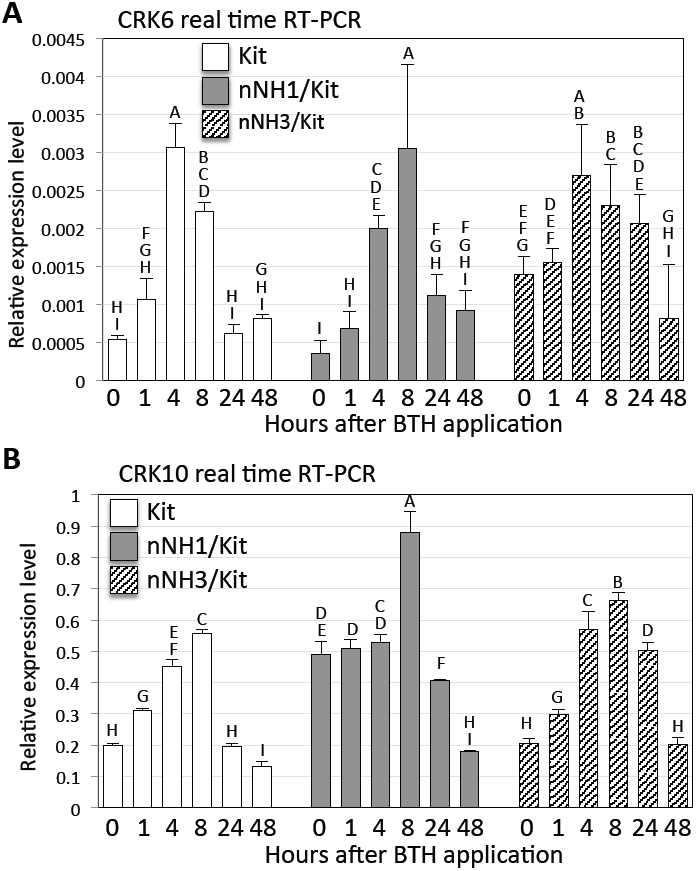
BTH treatment and elevated levels of *NH1* and *NH3* induce *CRK6* and *CRK10* expression. (A) *CRK6* RNA levels were determined by real time RT-PCR with tissues from Kit, nNH1 and nNH3 plants collected before BTH application and at 1, 4, 8, 24, and 48 hours after 1 mM BTH application. nNH1 are transgenic plants with an ectopic copy NH1 containing its native promoter driving its cDNA. nNH3 are transgenic plants with an ectopic NH3 containing its native promoter driving its cDNA. Both nNH1 and nNH3 are confirmed to express higher levels of NH1 and NH3, respectively. (B) *CRK10* expression levels determined by real time RT-PCR with the same RNA samples as described in (A). Each time point represents the average and standard deviation of three replicates. The letters above each bar represent the statistical groupings based on the 5% significance level.

### Silencing of *CRK6* and *CRK10* individually compromises NH1ox-mediated resistance

To further investigate the involvement of *CRK6* and *CRK10* in BTH-induced, NH1-mediated immunity to *Xoo*, we used the RNA interference (Ri) method to silence each of the two genes. We generated transgenic rice lines in the NH1ox background with constructs targeting each of *CRK6* and *CRK10* individually. T1 progeny were analyzed by inoculation with PXO99. The NH1ox-mediated resistance to PXO99 was found compromised in silencing lines. The presence of the *CRK10Ri* transgene was tightly correlated with susceptibility to *Xoo* (progeny of three lines shown in Supplemental Fig 1). The presence of the *CRK6Ri* transgene was also correlated with susceptibility (four lines presented in Supplemental Fig 2) although more variations in lesion length were observed for CRK6Ri. Independent lines with a 3:1 (PCR positive to negative) segregation ratio were selected based on PCR genotyping results. Three homozygous lines for CRK6Ri and CRK10Ri each were then selected after genotyping to confirm the presence of the transgene in all T2 progeny. More detailed *Xoo* inoculation analyses were carried out with homozygous lines as below.

The silencing of *CRK6* and *CRK10* in CRK6Ri and CRK10Ri lines were confirmed by real time RT-PCR (Supplemental Fig 3). *CRK6* RNA levels in CRK6Ri and CRK10Ri lines and in the NH1ox control were examined using a primer pair (G690-Q1a in Supplemental Fig 3) targeting a region specific to *CRK6* (see Methods). *CRK6* RNA levels in CRK6Ri lines showed an 80-85% reduction compared to the control. Unexpectedly, *CRK6* RNA levels were also reduced in CRK10Ri plants, even though *CRK6* and *CRK10* do not share high similarity at the nucleotide level (68% identity in the CRK10Ri target region). To confirm these results, another pair of primers (G690-Q1b) was used to repeat the real time RT-PCR and the results appeared similar. When *CRK10* RNA levels were assessed (with primer pair G700), the CRK10Ri lines showed a 75-90% reduction compared to the control. *CRK10* levels were only slightly affected in two out of the three CRK6Ri lines. Together, the RT-PCR results confirmed silencing of the *CRK6* and *CRK10* genes. Notably, *CRK6* was expressed at a level hundreds of times lower than those of *CRK10* and actin, which was used as the reference gene in the real time RT-PCRs.

Two homozygous lines each of CRK6Ri (#3 & #10) and CRK10Ri (#4 and #13) were inoculated with PXO99. As shown in Figure 7A, these CRK6Ri and CRK10Ri lines did not display lesion mimic necrotic spots whereas the NH1ox parent displayed typical necrotic spots (indicated by the white arrowheads) even in the absence of BTH treatment. These results indicate that both CRK6Ri and CRK10Ri suppressed the NH1ox-mediated necrotic development. Figure 7B shows two representative leaves from each line, two weeks after PXO99 inoculation. Figure 7C shows the development of lesions for each line over 12 days after inoculation. Consistently, these results indicate that silencing of *CRK6* or *CRK10* compromised NH1ox-mediated resistance. Bacterial growth curve analysis were carried out and the results are shown in Figure 7D. The results showed that CRK6Ri plants harbored 5-8 fold and CRK10Ri plants 11-12 fold more *Xoo* than the NH1ox plants at day 12, indicating that both CRK6Ri and CRK10Ri plants were more susceptible than the NH1ox parent. Notably, CRK10Ri plants were consistently more susceptible than CRK6Ri plants as measured by both lesion lengths and bacterial growth curve analyses. In summary, these results demonstrate that *CRK6* and *CRK10* are required for the BTH-induced, NH1-mediated immunity and their disruption contributes to the *snl1* phenotype.

**Figure 7.**
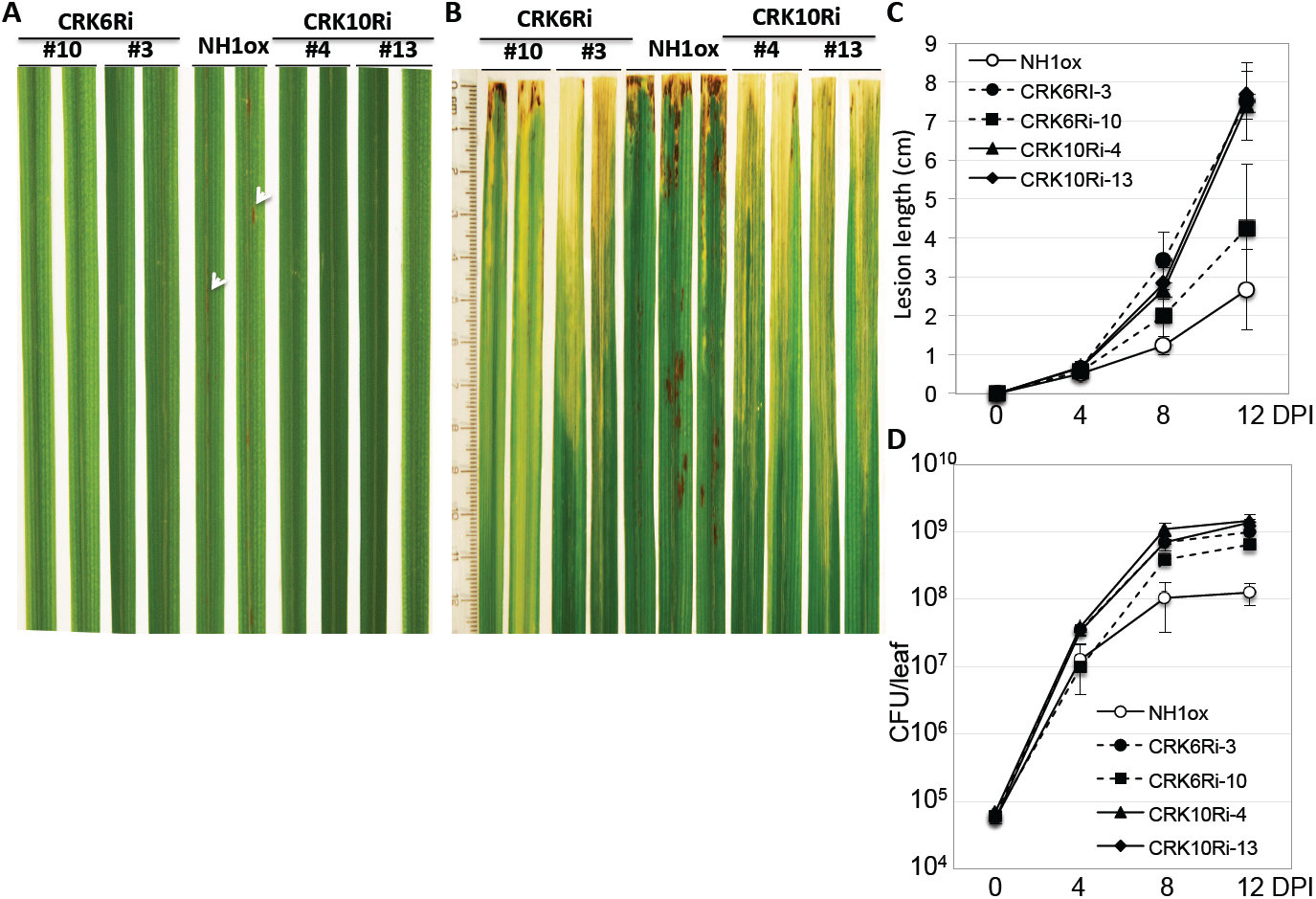
Silencing of *CRK6* or *CRK10* re-creates the *snl1* phenotypes. Two lines each of CRK6Ri (#3 and #10) and CRK10Ri (#4 and #13) are presented. Plants were treated with 1 mM BTH one day before *PXO99* inoculation. (A) Lesion mimic formation. Two representative leaves are shown for each line. The necrotic spots in the NH1ox parent are marked with white arrowheads. (B) *Xoo*-induced water-soaked disease lesions. Two representative leaves are displayed for each line. (C) Lesion length development over 12 days. Each time point represents the average and standard deviation of 6 leaves. (D) Bacterial growth curves. *Xoo* populations (colony forming unit/leaf) in inoculated leaves were determined for 12 days after *Xoo* inoculation (DPI). Each time point represents the average and standard deviation of three samples. Each sample contained two leaves. Statistical analysis of the day 12 bacterial population data reveals four significantly different groups: A=CRK10Ri-4, AB=CRK10Ri-13, BC=CRK6Ri-3, C=CRK6Ri-10, D=NH1ox.

### Inducible expression of *CRK10* enhances resistance to *Xoo*

The silencing experiments indicated that CRK6 and CRK10 are required for resistance to *Xoo* and suggested that overexpression of these genes might result in enhanced resistance. To test this hypothesis, we first tried to over-express *CRK6* (*Ubi-CRK6*) and *CRK10* (*Ubi-CRK10*) with the maize *ubiquitin-1* (*Ubi-1*) promoter in the rice Kitaake genetic background. We obtained 30 independently transformed Ubi-CRK6 lines and tested them for resistance to *Xoo*. None of these Ubi-CRK6 lines showed obvious enhanced resistance to *Xoo* (data not shown). Although we obtained equal number (>30) of transgenic, green calli for Ubi-CRK10, only a small number of these plants were able to regenerate, survive and grow in the greenhouse; most of these plants displayed lesion mimic necrotic spots and were dwarfed and unable to set seeds. These results suggested that over-expression of *CRK10* under the *Ubi-1* promoter leads to cell death.

As an alternative approach to assess the phenotypic effect of overexpression, we generated an inducible construct (GVG-CRK10) using the GVG (Gal4-VP16-Glucocorticoid receptor)-based, dexamethasone (DEX)-inducible expression system [41,42]. Approximately 10 healthy, independently transformed lines were obtained in the rice Kitaake genetic background. Our initial *Xoo* inoculation results of 5 of these lines indicated that most of the GVG-CRK10 lines displayed enhanced resistance to *Xoo* after DEX induction (Supplemental Figure 4). The enhanced resistance in the progeny co-segregated with the presence of the *GVG-CRK10* transgene. To further characterize these lines, we developed homozygous lines for GVG-CRK10-3, -21, and -32 (see Methods). We inoculated these lines with PXO99, (except for line #32 because the homozygous progeny of this line were dwarfed and unhealthy). Figure 8A shows two representative leaves for each line two weeks after inoculation. Figures 8B and 8C present the results of disease lesion length and bacterial growth curve analyses measured over 12 days after inoculation. In parallel, we also carried out real time RT-PCR on GVG-CRK10-3, -21, -32, and the Kit control after DEX application to assess the *CRK10* transcript levels. Figure 8D presents the real time RT-PCR results of plants at 0, 4, 24, and 44 hours after DEX induction. GVG-CRK10 line #21, which expressed higher levels of *CRK10* and was further induced by DEX application, showed high levels of resistance to PXO99 (Fig 8A, 8B, and 8C). GVG-CRK10 line #3 showed slightly higher, but statistically significant, level of *CRK10* at four hours after DEX induction (Fig 8D); consistently, it developed slightly shorter lesions at days 8 and 12 (Fig 8B) and harbored modestly lower levels of *Xoo* populations at days 4 and 8 (Fig 8C) than Kit. In contrast, Kit was highly susceptible to *Xoo*. Notably, the difference between the *Xoo* populations in GVG-CRK10-21 and Kit at day 12 was nearly 1,000-fold. These results confirm that elevated levels of *CRK10* expression lead to extremely high levels of resistance to *Xoo*. GVG-CRK10 line #32 expressed high levels of *CRK10* RNA, which might have caused its impaired development.

**Figure 8.**
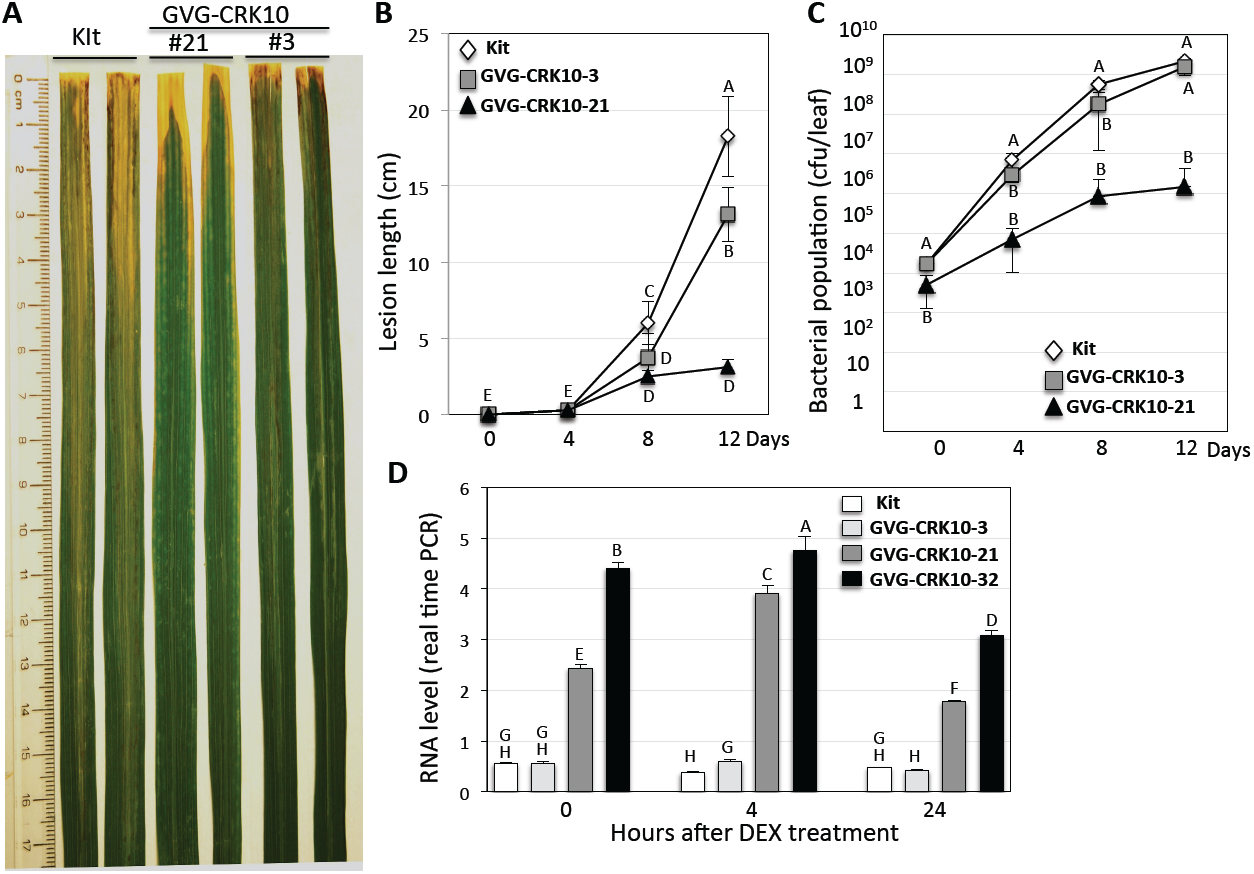
Inducible over-expression of *CRK10* enhances resistance to *Xoo*. (A) Two leaves (2 weeks after *PXO99* inoculation) each of GVG-CRK10 lines (#3 and #10) and the Kitaake control are presented. (B) Lesion development and (C) bacterial growth curves were determined over 12 days after *PXO99* inoculation. Each time point represents the average and standard deviation of four leaves. The letter next to each time point represents the statistical grouping based on the 5% significance level. (D) *CRK10* RNA levels in GVG-CRK10 lines (#3, #21, and #32) and the Kitaake control. Real time RT-PCRs were done with RNA extracted from leaves collected before DEX application (0 hours) and at 4, 24, and 44 hours after 100 µM DEX application. The letter above each bar represents the statistical grouping based on the 5% significance level.

After DEX treatment, many of the GVG-CRK10 lines, such as lines #21 and #32, exhibited a severe lesion mimic phenotype — a phenotype similar to that of the NH1ox parental line. These results indicate that the expression level of *CRK10* is critical for disease resistance and lesion mimic formation.

### The Arabidopsis proteins AtCRK6, AtCRK8, and AtCRK10, are similar to rice CRK6 and CRK10

In a BlastP query of Arabidopsis proteins with CRK6 and CRK10, AtCRK6, AtCRK8, and AtCRK10 were identified as the most similar proteins. All three Arabidopsis proteins share ca. 43% identity with rice CRK6 and CRK10. When a phylogenetic tree was constructed for available Arabidopsis CRK protein sequences, AtCRK6, AtCRK8, and AtCRK7 form a clade, and together with AtCRK10 and AtCRK15 form a larger clade (Supplemental Figure 5). This clade is distinct from the clades containing AtCRK5, AtCRK13, and AtCRK45, which have been implicated in immune responses [43–45]. Similar to rice CRK6 and CRK10, which are clustered on rice chromosome 7, AtCRK6, AtCRK8, and AtCRK10 are clustered at the same locus on Arabidopsis chromosome 4. Based on this analysis, we hypothesize that AtCRK6, AtCRK8, and AtCRK10 are orthologous to rice CRK6 and CRK10, and thus may likely also regulate the plant immune response. None of these proteins have yet been characterized.

### Discussion

The *snl1* mutation described here was caused by an 88-kb deletion, which contains several *CRK* genes including the previously uncharacterized genes, *CRK6* and *CRK10*. Complementation of *snl1* and gene silencing experiments showed that *CRK6* and *CRK10* are responsible for the *snl1* phenotypes, including suppression of lesion mimic formation and immunity to *Xoo*. These results indicate that rice CRK6 and CRK10 are required for the rice innate immunity induced by the plant defense activator BTH, whose function is mediated by the NPR1-like proteins NH1 and NH3.

Although altered expression of *CRK* genes has been observed in several datasets in response to biotic and abiotic stress [43–46], the biological functions of CRK proteins have not been well-characterized. In previous studies, controlled over-expression of *AtCRK5* and *AtCRK13* led to hypersensitive response and cell death, and activation of defense genes [43,44]. *AtCRK45* was recently suggested to positively regulate resistance to *Pseudomonas syringae* pv. tomato DC3000 [45]. Two other reports suggested that *AtCRK20* and *HvCRK1* genes negatively regulate immune responses in Arabidopsis and barley, respectively [47,48]. To our knowledge, little genetic evidence has been reported on the biological function of *CRK* genes. Here we provide direct evidence that the previously uncharacterized proteins CRK6 and CRK10 mediate the NH1-mediated immune response.

Silencing of *CRK6* partially re-creates the *snl1* phenotype. In contrast, co-silencing of *CRK10* and *CKR6* in the CRK10Ri lines fully re-creates the *snl1* phenotype. The more severe phenotype of the CRK10Ri plants may be partly due to the fact that *CRK6* expression is down-regulated in the CRK10Ri plants. Although *CRK6* and *CRK10* share only 68% sequence identity at the nucleotide level in the regions used for RNAi, there is a stretch of 19 identical nucleotides shared between these two sequences. This stretch of nucleotides may be responsible for the co-silencing of *CRK6* observed in the CRK10Ri lines. However, even though this stretch of nucleotides was present in the CRK6Ri construct, it did not affect *CRK10* expression. Thus, we do not rule out the possibility that *CRK6* was co-silenced via a different mechanism.

The CRK6 and CRK10 proteins contain a presumed extracellular domain that is rich in cysteine residues. Cysteine residues commonly form disulfide bonds under oxidative conditions and these disulfide-bonds are disrupted in response to a rise in redox levels, causing a conformational change in these proteins. Thus, the cysteine-rich domains may serve as sensors of the cellular redox status. In support of this hypothesis, overexpression of rice CRK10 or Arabidopsis CRK13 [44] enhances cell death in rice and Arabidopsis, respectively. These observations support a model where high levels of CRK proteins can trigger cell death in the absence of pathogen infection in response to an endogenous activation signal that affects cellular redox status.

It is well known that NH1 interacts with rice TGA transcription factors. We have also shown that NH3 also interact with the same set of TGA transcription factors (Chern and Ronald, submitted). NH1 also serves as a transcriptional co-activator to activate downstream genes through interaction with a TGA protein, which is anchored to a promoter in a protoplast transient assay [37]. Based on the structural and functional similarities of NH3 and NH1, we hypothesize that NH3 is likely to function in the same manner as NH1.

The *CRK6* promoter contains a cognate sequence of TGA binding sites (TGACG) located 164 bp upstream of the putative TATA box. The *CRK10* promoter contains two such sequences: one (TGACG in reverse orientation) 381 bp and the other (TGACGT) 800 bp upstream from the putative TATA box. Thus, *CRK6* and *CRK10* promoters are likely targets of rice TGA proteins and may also be regulated by NH1 or NH3. We have observed that the *CRK6* transcript level is responsive to elevated level of *NH3* transcripts. In contrast, the *CRK10* transcript level is elevated in response to higher level of *NH1* transcripts. It is not known why higher *NH1* levels failed to increase *CRK6* expression and higher *NH3* levels failed to elevate *CRK10* expression. (Fig 6).

Figure 9 summarizes our current model for how the plant defense activator BTH activates defense responses via NH1/NH3 and CRK6/CRK10. In this model, the NH1 and NH3 proteins act as receptors to perceive and respond to the activators. Activated NH1 and NH3 proteins are translocated to the nucleus where they interact with transcription factors, such as TGA proteins, and function as transcriptional co-activators to turn on expression of early immediate downstream genes, such as *CRK6* and *CRK10*. CRK6 and CRK10 proteins are localized to membranes where they possibly interact with other receptor kinases and signaling components to trigger downstream defense signaling, leading to activation of defense responses and immunity.

**Figure 9.**
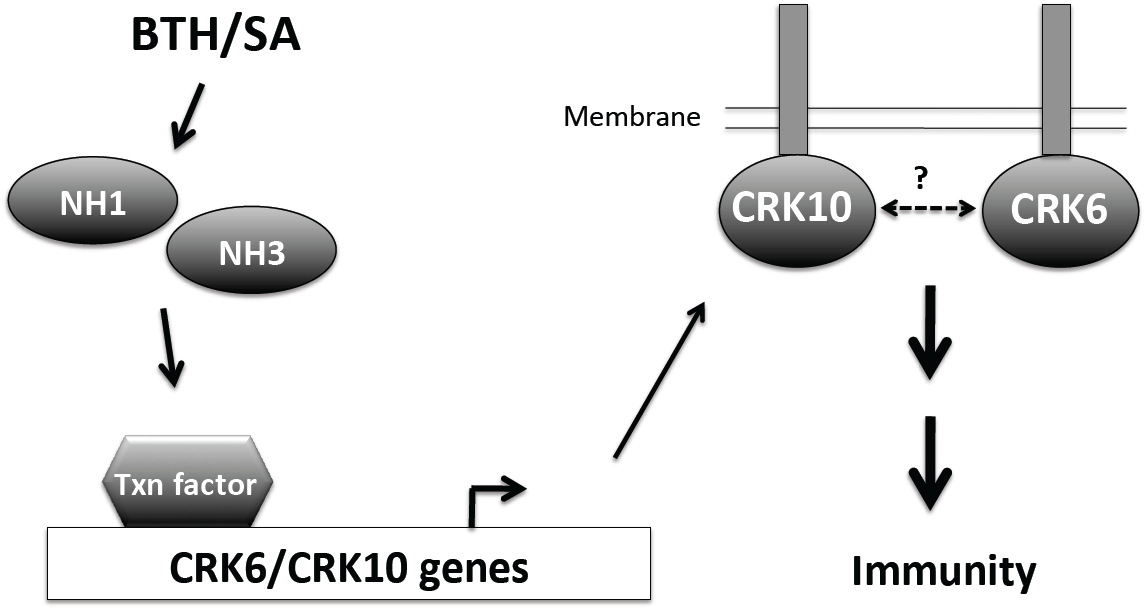
A model for NH1/NH3 and CRK6/CRK10 function. NH1 and NH3 may directly perceive plant defense activators BTH or the endogenous hormone SA. Activated NH1 and NH3 proteins are then anchored by transcription (Txn) factors, such as TGA proteins, to the promoters of *CRK6/CRK10* genes to induce their expression. The CRK6 and CRK10 proteins are localized to membranes where they may interact with each other and with other receptor kinases and signaling components to transduce the signaling, leading to immune response.

It remains possible that transcription factors other than TGA factors may also interact with NH1 and/or NH3 and regulate *CRK6* and *CRK10* transcription. In support of this hypothesis, a wheat NPR1-like protein was reported to interact with a wheat WRKY transcription factor [49]. In addition, two novel transcription factors, in addition to the characterized TGA factors, have recently been reported to interact with Arabidopsis NPR1 and mediate its function (Xinnian Dong, personal communication).

## Materials and Methods

### Plant materials and screening

The rice mutant population in the NH1ox-54 genetic background (in the rice LG variety carrying the *Ubi-NH1* gene) was generated by irradiation with fast neutron at 20 Grey as previously described [40]. The irradiated rice seeds were grown in a rice field at the University of California, Davis. Seeds (F2) from the first generation were harvested in groups of ten lines. Approximately 150 seeds from each group were planted in order to screen on F2 plants in the UC Davis rice field. BTH was applied twice to the plants at a concentration of 10 mM to ensure adequate application of the inducer and to minimize escapes. Seeds of candidate mutants were harvested and re-tested in a green house.

### Plant growth and *Xoo* inoculation

For *Xoo* inoculation, plants were grown in a greenhouse till approximately 6 weeks old. Plants were moved to a growth chamber, set at 26°C with 80% humility, in a control environmental facility. Inoculation with *Xoo* strain PXO99 was carried out with the scissor-dip method as described [50]. PXO99 was grown on a PSA plate with cephalexin (20 µg/mL) for two to three days. PXO99 cells were recovered with a Q-tip from the plate and re-suspended in sterile water. The absorbance (600 nm) of *Xoo* concentration used for inoculation was at OD=0.5.

### Comparative Genome Hybridization

Comparative genome hybridization was carried out at the Roche NimbleGen facility (Madison, WI) using the Roche NimbleGen rice whole genome tiling array, which contains 2.1 million probes of 50-70mer oligonucleotides [40]. Rice genomic DNA was extracted from the *snl1* mutant and from the NH1ox parent separately, with a method using the CTAB extraction buffer, and further purified with a plant genomic DNA purification kit (DNeasy Plant Mini Kit) from Qiagen (Maryland, USA).

### Cloning of individual genes on chromosome 7 for test to complement the *snl1* mutant

A Qiagen Long Range PCR kit was used for amplification of each gene including the promoter, the coding region, and the 3’ sequence, from chromosome 7. Amplification of g35580 used primers G580-1 (CACCTATTGT TTGAGCTACA TGTGGACATCA) and G580-3 (ACCAGTGAGT ACACTACTCT ATTC), g35600 used primers G600-1 (CACCTGCGAC TTCTCCGTTA GCTGTCGG) and G600-2 (CTTCACCATC CGCCACTAAA AAGC), g35610 used primers G610-1 (CACCGTGACG TGTCTGTCTC ACTG) and G610-3 (GGAATAATAA CAAACAGGCC TAACACCC), g35620 used primers G620-1a (CACCTCATGG AGGAGTCGCG GA) and G620-2a (TCAGGACGCT GGGGTGAAAG), g35630 used primers G630-1 (CACCTAAGCA TGTCTAAGCA ATTCCTAGTCCA) and G630-2 (GATCTTCAGC CGTGATTCTT CATGG), g35640 used primers G640-1 (CACCGTATTT CACCTGTAAA CTGCGAGATG) and G640-2 (ACACAAGATT GGCTACATGG GCATCGAGA), g35650 used primers G650-1 (CACCGAATTT TGCTCCTTTC TATATCAGCT TCAATGG) and G650-2 (CCTTTATTGT GCGCACAAAT ACAGGT), g35660 used primers G660-1 (CACCTAGGCA ATAGAGAATC GGATAGTGA) and G660-2 (TTCGGGCAGT GTAGAGTAGA TGTTG), g35680 used primers G680-1 (CACCCGGTCC AGAAATCCGG ATTTCCT) and G680-3 (GACCGATACC AGTACCACTC GG), g35690 used primers G690-1 (CACCTCCCTA TTCTCAGTTC TAGAACCAAG CA) and G690-3 (GTGCGTTAAA AAGTTCAAAG TCGTATCTCC GGT), and g35700 used primers G700-1 (CACCTCCCTT CCATGCTTCT CAAACC) and G700-2 (GGTCAGGATC TTGCTTGTAG GGA).

Amplification of g35600, g35610, g35640, g35650, g35660, g35690, g35700 used Liaogeng (LG) genomic DNA as the PCR template. Due to difficulty in PCR reaction, amplification of the remaining genes used PAC clone P0458H05 as template. PCR products were cloned into the pCR8/GW/TOPO vector (Invitrogen) and confirmed by sequencing. Each gene was sub-cloned into the C4300 vector by the Gateway recombination. The resulting constructs were used to transform the *snl1* mutant using mannose selection.

### Plasmid construction for gene silencing and over-expression

To generate an RNAi construct targeting *CRK6* (Os07g35690), we used primers G690-SiRI (TTTGAATTCA CCAGGTCAAC CTCGACCTC) and G690-SiBam (TTGGATCCAG TTGCCCGATC ACCGTCGAGA) to amplify a 500-bp fragment from the 5’-end of *CRK6*. This fragment was digested with EcoRI and BamHI and cloned into a plasmid (pENTR/L16) modified from the pENTR/D vector to contain multiple cloning sites. The clone was confirmed by sequencing. The fragment was excised with EcoRI and BamHI and sub-cloned into pBluescript II SK-, pre-cut with BamHI and phosphatase-treated, jointly with the *Xa21* intron (precut with EcoRI). The resulting clone (dsG690/SK) contained two pieces of the *CRK6* fragment head-to head with the *Xa21* intron in between to serve as a spacer to stabilize the clone in bacteria. The dsG690 insert was excised with BamHI and sub-cloned back to the pENTR/L16 vector using the BamHI site. The resulting construct dsG690/L16 was used to recombine with a Gateway compatible Ubi-C4300 binary vector (Ubi-C4300/GA) and resulted in construct Ubi-dsG690/C4300. This construct was used to transform the NH1ox-11 rice line, which is hygromycin resistant, using the mannose selection generating CRK6Ri lines in the NH1ox genetic background.

To generate an RNAi construct targeting *CRK10* (Os07g35700), we used primers G700-SiRI (TTTGAATTCA CTACACGGAG CACGGCACG) and G700-SiBam (TTTGGATCCA TGTCTGGCGT GCACTGC) to amplify a 500-bp fragment from the 5’-end of *CRK10*. The PCR product was processed the same way as the *CRK6* fragment for generating the end product Ubi-dsG700/C4300 construct. This construct was also used to transform the NH1ox-11 line, generating CRK10Ri lines in the NH1ox background.

A full-length, 2-kb *CRK10* cDNA was amplified with primers G700-3 (CACCATGTCC ATGGCCTGCT ACTACC) and G700-8 (TCTTGAGTTG TGTGGGTTC) using a cDNA pool from Nipponbare. The PCR product was cloned into the pENTR/D vector and confirmed by sequencing. The cDNA was sub-cloned to the Ubi-C1300 binary vector through Gateway recombination, generating Ubi-G700/C1300. To generate an inducible *CRK10* construct in the GVG-DEX system to over-express *CRK10*, the same *CRK10* cDNA was sub-cloned into vector TA7002/GA by Gateway recombination, creating binary construct GVG-G700. Genotyping of the GVG-G700 construct in transgenic lines used primers Hyg-3 (TCCACTATCG GCGAGTACTT CTACACA) and Hyg-4 (CACTGGCAAA CTGTGATGGA CGAC), targeting the hygromycin selection marker. Lines carrying a single insertion were selected based on a 3;1 (PCR positive to negative) segregation ratio. Homozygous lines were then obtained via identifying lines where their progeny were all PCR positive by genotyping approximately 18 progeny plants.

Due to the extremely low expression level of *CRK6*, we were unable to clone a full-length *CRK6* cDNA. A genomic clone containing full-length *CRK6* was amplified from PAC clone P0458H05 and used for over-expression. This 3-kb genomic DNA was amplified with primers G690-3b (GGATTCATCT CGGCACAAGC TCTGTG) and G690-4c (CACCATTACC ATCGCCAGCA CA), cloned into the pENTR/D vector, and confirmed by sequencing. The *CRK6* insert was sub-cloned into Ubi-C1300 by recombination, generating Ubi-G690/C1300. These binary constructs were used to transform Kit rice variety.

### Real time quantitative RT-PCR

Total RNA was extracted using the Trizol reagent following the manufacturer’s instruction (Invitrogen). Extracted total RNA was precipitated with isopropanol and rinsed with 70% ethanol and dried down. RNA was then resuspended in 90 µl of RNase-free H2O. DNase I was added to remove residual DNA in a volume of 100 µl. The DNase I enzyme was removed by treating the RNA sample with 300 µl of the Trizol reagent and then 100 µl of chloroform. The supernatant was passed through a RNA purification spin-column (NucleoBond) to purify the RNA following the manufacturer’s instruction. Three to five µg of total RNA each sample was used to synthesize cDNA for real time RT-PCR.

To assess the expression level of *CRK6*, primers G690-Q1a (CCAAAGAATT CAGCGGGAGG) and G690-Q2 (GTCGCCGATG GCGAAGGC) or primers G690-Q1b (TCGACGGTGA TCGGGCAACT) and G690-Q2 (both pairs targeting the C-terminus of its putative extracellular region) were used in real time RT-PCR. These primers were determined to be specific to the *CRK6* gene. For *CRK10*, primers G700-RT3 (TTTGGCTCCT ACGGTTCTGAC) and G700-RT5 (CACAGAGTAG CCCAATGTGGA) (targeting the C-terminus of *CRK10* reading frame) were used for real time RT-PCR.

### Genotype determination of plants carrying the *snl1* deletion and RNAi constructs

Genotype determination of plants carrying the 88-kb deletion was mainly carry out with primers targeting genes *CRK6*, *CRK10*, or Os07g35610. *CRK6* genotyping used primers G690-RT1 (GATAGTGGGC AAGATGTTGA TCTC) and G690-RT2 (TGGATAGCGT TCTGAATACG GA). *CRK10* genotyping used primers G700-RT3 (TTTGGCTCCT ACGGTTCTGAC) and G700-RT4 (TGCACCTTAG CTAGCAGTAG CA). Os07g35610 genotyping used primers G610-10 (CACCATGATG GTGAAGAAAA AAACCTCTTG GAGC) and G610-2 (GCTACTAAGT TGGTGGATTA CCTAGC).

Genotype determination of CRK6Ri plants used primers G690-SiRI (listed above) and Ubi-1 (TGATATACTT GGATGATGGCA) primers; genotyping of CRK10Ri plants used G700-SiRI (listed above) and Ubi-1 primers. For genotyping of RNAi plants, genomic DNA was digested with EcoRI before PCR amplification.

### BTH and DEX applications to plants

BTH was applied to rice leaves by foliar spray. For the mutant screen in the rice field, BTH was applied at 10 mM in the form of Actigard. For other applications, BTH was applied in a greenhouse at 1 mM in the form of Actigard. DEX was dissolved to 100 µM in 0.05% Tween 20 and applied by foliar spray.

### Statistical analysis

Statistical analysis was carried out using the JMP Pro 10 statistics program.

## Acknowledgements

We thank the National Institute of Agrobiological Sciences (NIAS) in Japan for the generous gift of PAC clone P0458H05. We thank Benjamin Schwessinger for helpful advice and Benjamin Schwessinger and Weiguo Zhang for careful editing of the manuscript.

---

**Supplemental Figure 1.**
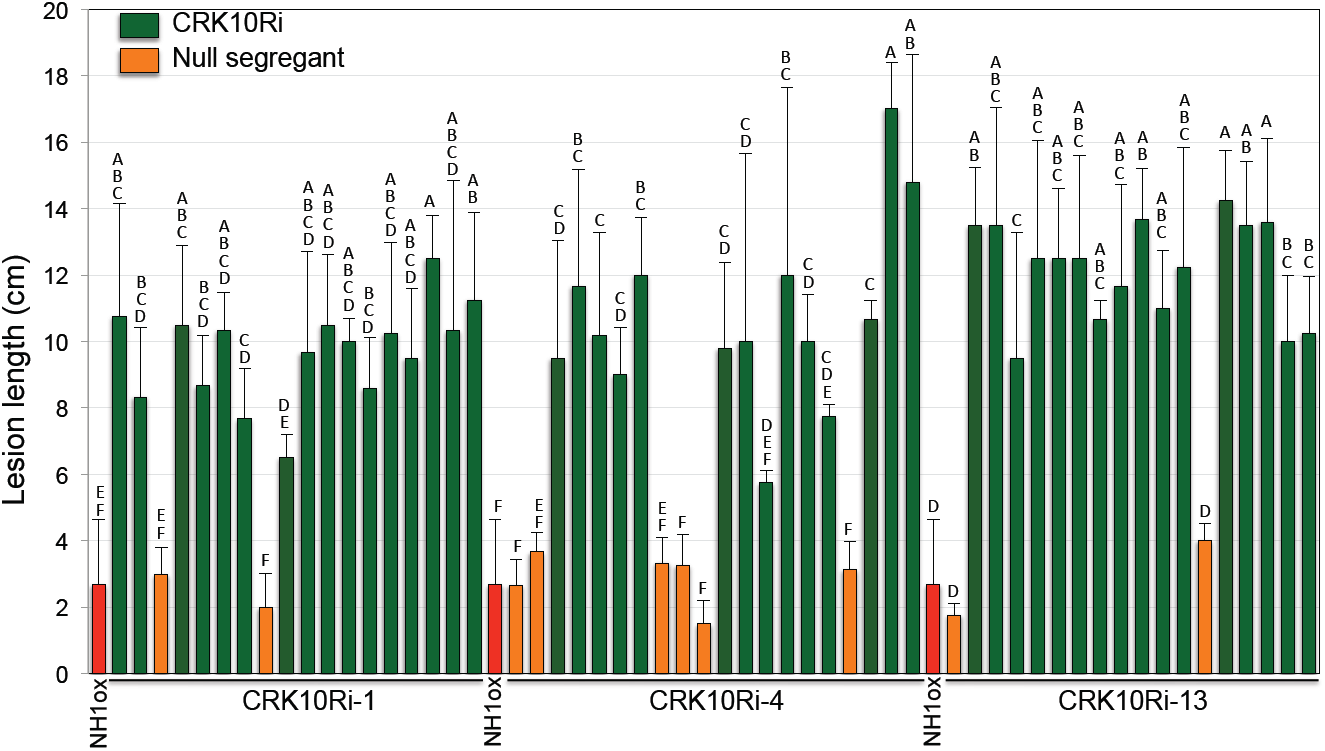
The CRK10Ri construct cosegregates with enhanced susceptibility in progeny. Segregating progeny were genotyped for the presence of the CRK10Ri transgene. Those containing the transgene colored in green and the null segregants colored in orange. Progeny plants and the NH1ox parent were inoculated with *PXO99*. Lesion lengths were measured two weeks after inoculation. The inoculation results of three lines are presented. Each bar represents the average lesion length and standard deviation of all inoculated leaves from one plant. The letters above each bar show the statistical groupings using the student T-test on each pair based on the 5% significance level within the progeny of each line plus control.

**Supplemental Figure 2.**
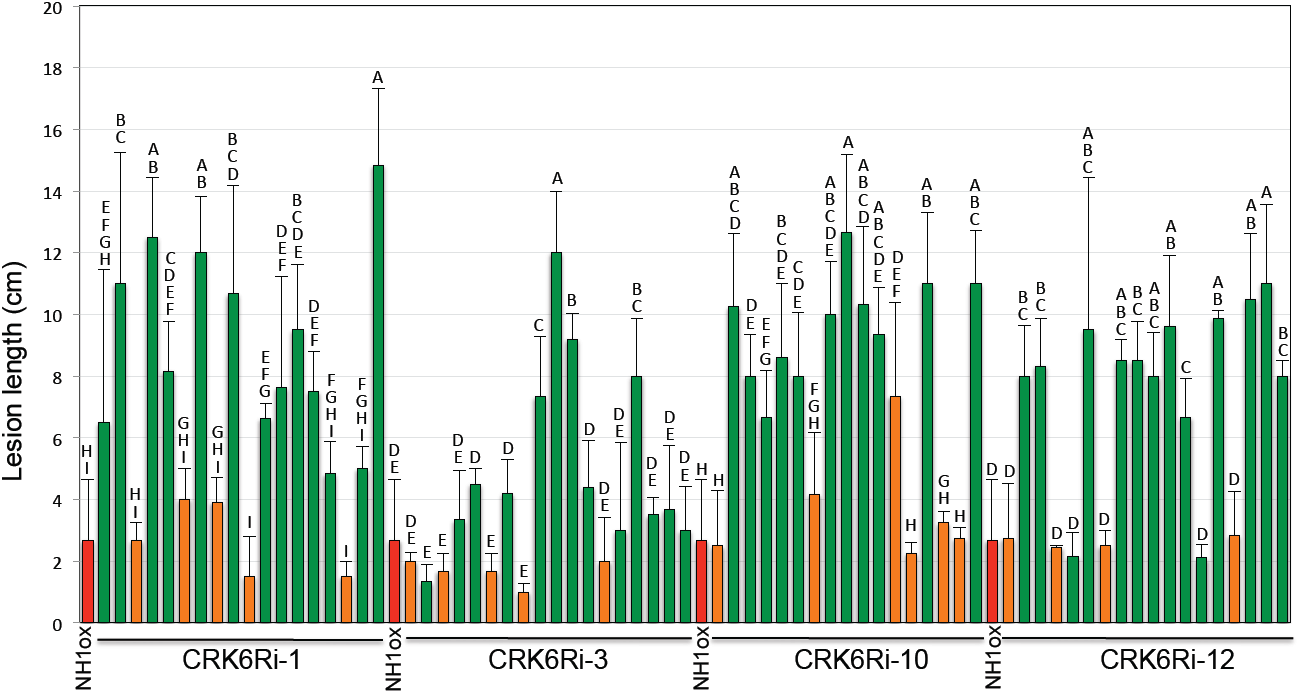
The CRK6Ri construct cosegregates with enhanced susceptibility. Segregating progeny were genotyped for the presence of the CRK6Ri transgene. Those containing the transgene colored in green and the null segregants colored in orange. Progeny plants were inoculated with *PXO99* together with the NH1ox parent. Lesion lengths were measured two weeks after inoculation. The inoculation results of four lines are presented. Each bar represents the average lesion length and standard deviation of all inoculated leaves from one plant. The letters above each bar show the statistical groupings using the student T-test on each pair based on the 5% significance level within the progeny of each line plus control.

**Supplemental Figure 3.**
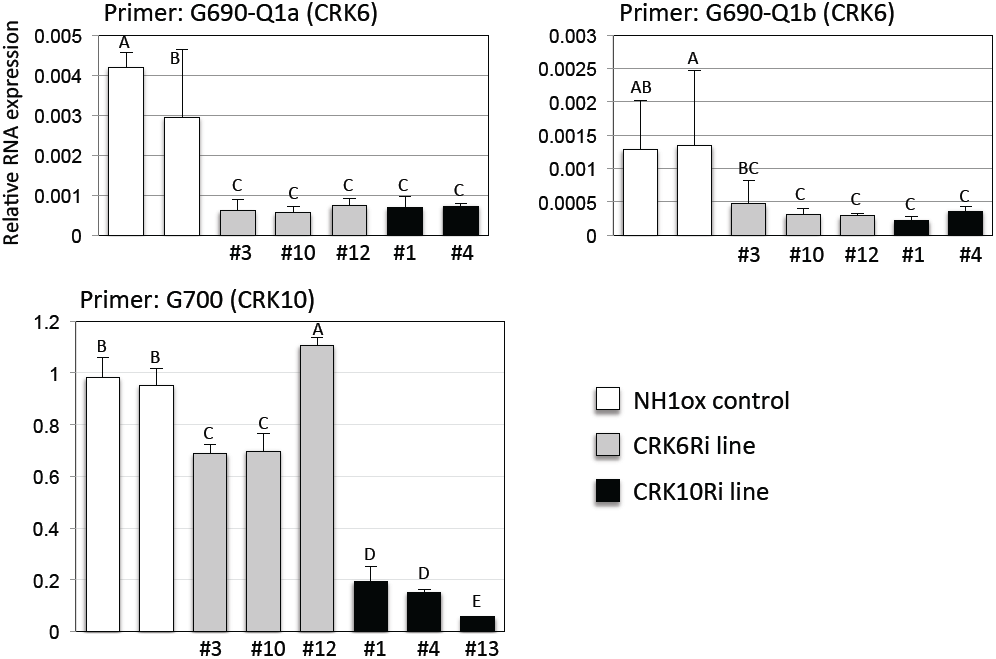
*CRK6* and *CRK10* are silenced in the CRK6Ri and CRK10Ri lines, respectively. RNA was extracted from independent CRK6Ri and CRK10Ri lines as labeled under each bar. *CRK6* RNA levels were determined by running real time RT-PCR with primers G690-Q1a and G690-Q2, which are specific to the *CRK6* gene. Real time RT-PCRs were also carried out with G690-Q1b and G690-Q2 to confirm the above PCR results. *CRK10* RNA levels were assessed with primers G700-RT3 and G700-RT5, which are specific to the *CRK10* gene. Each bar represents the average and standard deviation of three replicates. The letters above each bar show the statistical groupings using the student T-test on each pair based on the 5% significance level.

**Supplemental Figure 4.**
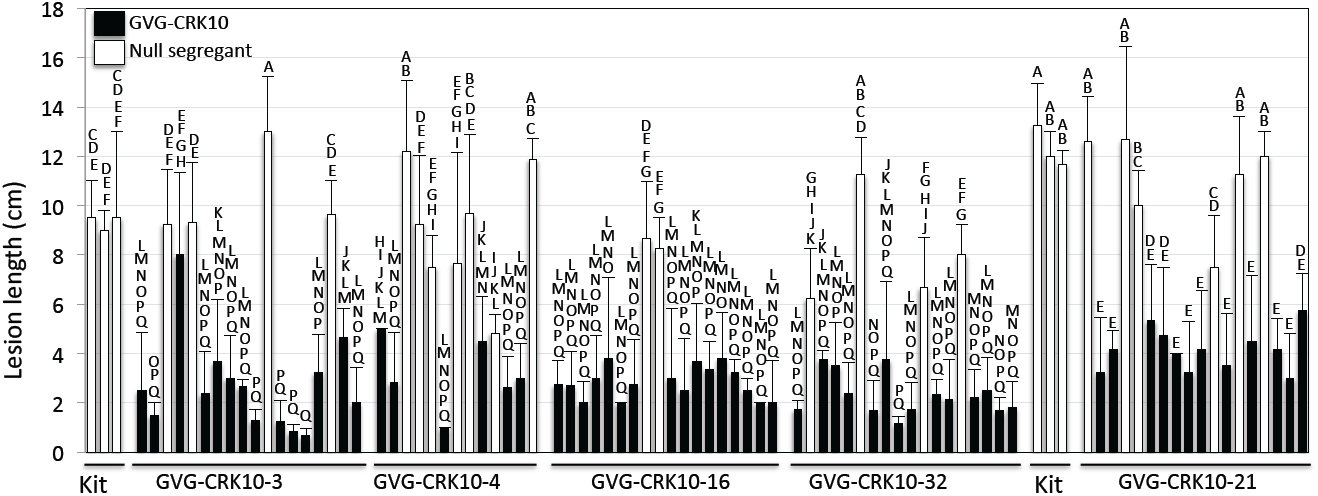
The GVG-CRK10 construct cosegregates with enhanced resistance. Segregating progeny were genotyped for the presence of the GVG-CRK10 transgene. Those containing the transgene presented in filled bars and the null segregants in open bars. Progeny plants were inoculated with *PXO99* together with the Kit control after DEX induction. Lesion lengths were measured two weeks after inoculation. The results of four lines from one inoculation and another line from another inoculation are presented. Each bar represents the average lesion length and standard deviation of all inoculated leaves from one plant. The letters above each bar show the statistical groupings using the student T-test on each pair based on the 5% significance level. Progeny of lines #3, #4, #16, and #32 are compared together with the Kit control. Progeny of line #21 were compared with its own Kit control separately due to the different inoculation time. The letters above each bar show the statistical groupings using the student T-test on each pair based on the 5% significance level within the progeny of each line plus control.

**Supplemental Figure 5.**
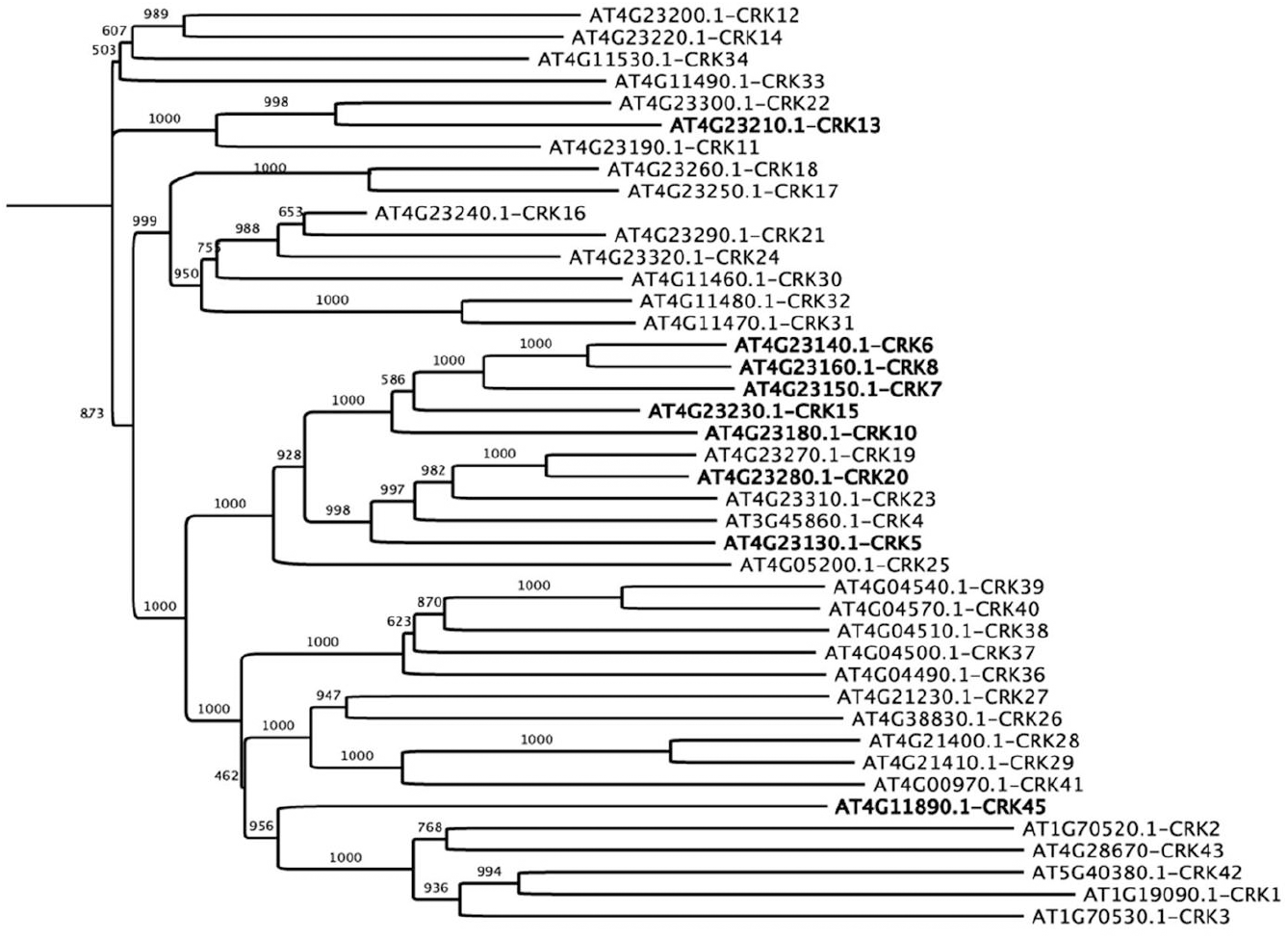
A phylogenetic tree of Arabidopsis CRK proteins. Arabidopsis CRK protein sequences were retrieved from the NCBI protein database and used to generate the phylogenetic tree using Clustal X based on the neighbor-joining method. AtCRK6, AtCRK8, and AtCRK10, which are clustered at the same locus on chromosome 4, share the highest identities and similarities with rice CRK6 and CRK10.

